# Auditory responses in the ventral tegmental area of awake, freely moving mice

**DOI:** 10.64898/2026.03.17.712298

**Authors:** Samira Souffi, Israel Nelken

## Abstract

The ventral tegmental area (VTA) is a key region in the reward system of the vertebrate brain, yet its role in sensory processing remains largely unexplored. Here, we use fiber photometry in awake freely-moving mice to investigate how the VTA represents auditory stimuli. We compared VTA responses to those in the inferior colliculus (IC) recorded with the same technique. Neural activity in the VTA exhibited robust responses to a wide variety of auditory stimuli, including broadband noise, pure tones, and music stimuli. We identified a subset of short-latency trials, comparable to those observed in the IC, with response durations that were longer than in the IC. VTA responses to complex sounds showed minimal envelope tracking and low temporal reproducibility, resulting in poor discriminability. This study positions the VTA as a potentially active player in shaping the perception of sound.

## Introduction

Emerging evidence suggests that neuromodulatory nuclei play a significant role in sensory processing (Engelhard et al., 2019; Glennon et al., 2023). In humans, fMRI studies have shown that rewarding auditory stimuli, such as music that evokes measurable pleasure, modulate the ventral tegmental area (VTA) and the nucleus accumbens (NAc), key components of the brain’s reward system (Blood and Zatorre, 2001; Menon & Levitin, 2005; Salimpoor et al., 2011, 2013). Notably, these studies revealed strong correlations between dopamine release and the activity of the NAc, suggesting that rewarding auditory experiences engage the VTA and cause dopamine-release. In addition to reward-related sound activity, other neuromodulatory systems also respond to auditory stimuli. Neurons in the locus coeruleus and the nucleus basalis, which are respectively the primary sources of norepinephrine and acetylcholine in the central nervous system, respond to auditory stimuli (Rasmussen et al., 1986; Hervé-Minvielle and Sara, 1995; Maho et al., 1995), and the dorsal raphe nucleus, source of serotonin, exhibits responses to auditory as well as visual inputs (Heym et al., 1982; Rasmussen et al., 1986). However, the way sensory information is represented in the neuromodulatory systems is poorly understood.

The VTA is part of the brain’s reward system, playing an essential role in processing motivational states. Dopaminergic neurons in the VTA are best-known for their activation by unexpected rewards and by cues that predict reward (Shultz et al., 1997). However, the VTA is involved not only in reward prediction and reinforcement but also in modulating sensory processing (Bao et al., 2001; Lou et al., 2014; Brunk et al., 2019; Herpers et al., 2022). In addition, electrophysiological recordings in awake, freely moving cats have shown that VTA dopaminergic neurons respond to unconditioned sensory stimuli, including auditory and visual inputs (Horvitz et al., 1997). More recently, Wei and colleagues (2024) conducted extracellular recordings in head-fixed awake naive mice and found significant VTA auditory responses to broadband white noise.

Here, we characterize neural responses in the VTA to auditory stimuli using a rich set of sounds, ranging from pure tones to complex sounds. Using fiber photometry, we recorded calcium activity in the VTA of awake freely-moving mice using a fast, small-molecule calcium indicator (Oregon Green BAPTA-1, OGB), in order to study the dynamics of auditory responses to envelope fluctuations at rates that are relevant for natural sound processing. We also recorded under the same conditions in the inferior colliculus, which served as a control region for auditory-evoked responses. We observed auditory-evoked responses in the VTA to all sound stimuli, although the VTA carried only coarse information about them.

## Materials and Methods

### Animals

Eight- to nine-week-old male mice (n = 17, C57BL/6JOLAHSD, Envigo, Israel) were used in this study. Male mice were selected to reduce variability related to the estrous cycle in females. All mice were kept on a 12-hour light-dark cycle in an animal facility with free access to food and water. Mice were housed in a group of same-sex littermates. All experimental procedures, surgeries and care of laboratory animals used in this study were approved by the Hebrew University Institutional Animal Care and Use Committee (IACUC). The Hebrew University of Jerusalem is an AAALAC-accredited institution.

### Surgical procedures

The mouse was anesthetized in an induction box (4% sevoflurane in oxygen, 0.1 L/min). After 10 minutes in the box, the animal was carefully transferred to a stereotaxic frame (Kopf Instruments, Tujunga, CA, USA) with its nose placed in a mask through which sevoflurane and oxygen mixture was delivered. Ophthalmic ointment was applied to prevent the eyes from drying. The animal was placed on a heating pad, and its body temperature was measured with a rectal thermometer and kept around 37.5 degrees. The depth of anesthesia was validated by pinching the tail. Lidocaine (0.1 mL, 5 mg/mL) was subcutaneously injected into the scalp overlying the skull as a local analgesic agent. In addition, meloxicam (10 mg/kg) was administered subcutaneously for systemic analgesia before surgery. The skin and muscles overlying the skull were removed and a short metal bar was glued along the sagittal suture of the skull with dental cement. The metal bar was then attached to a holder for the rest of the procedure. The muscles were gently retracted to reach the parietal bone for the VTA or the occipital bone for the IC. For VTA, we used lateral-medial (LM): -0.5 mm, rostral-caudal (RC): -3.2 mm, dorsal-ventral (DV): 4.7 mm, relative to Bregma; for IC, lateral-medial (LM): 1 mm, rostral-caudal (RC): 5 mm, dorsal-ventral (DV): 0.5 mm. These coordinates were based on the Paxinos and Franklin mouse brain atlas (Paxinos and Franklin, 2001). An opening with a radius of about 3 mm was made with a fine drill while cooling the surface with saline to avoid injury to the brain.

A glass micropipette filled with a solution containing the fluorescent dye OGB-1 AM (Invitrogen, Thermo Fisher Scientific, USA; see below for details) was progressively lowered using a micromanipulator (PatchPad, Scientifica, UK) into the brain. The dye was injected using air pressure of ∼10 PSI for ∼1 minute per site (PV830 Pneumatic Picopump, World Precision Instruments, USA). For the VTA, two injections were made along the dorsal–ventral axis: the first located 300 µm above the target coordinates, and the second at the exact target coordinates, to fully cover the region of interest. For the IC, a single injection was made at the target coordinates.

Approximately 30 min after dye injection, a single 0.2 mm optical fiber (Thorlabs, Inc., USA; Numerical Aperture 0.50; with a 1.25 mm ceramic ferrule and cut to a length of 6 mm) was implanted with the same micromanipulator into the stained region to the depth providing maximal fluorescence intensity, typically at about 100–300 μm above the injection site. Before implantation, the tip of the fiber was coated with a lipophilic red fluorescent dye (DiI; Invitrogen), for visualizing the fiber track after the experiment.

The optical fiber was positioned vertically, perpendicular to the brain surface, and temporarily secured using a custom connector that wrapped around a metal bar and was tightened in place with a screw. The fiber was then permanently affixed to the skull with a thin layer of dental cement (C&B Metabond, Parkell), followed by a thicker layer of acrylic dental cement.

All recordings were performed inside a sound-proof chamber (IAC 1202). Once the dental cement had fully cured, the mice were transferred to a 45 × 45 × 45 cm open field arena with access to wet food. Imaging was performed once the animals exhibited clear signs of wakefulness—such as exploratory movements, normal posture, and responsive behavior (e.g., whisking, grooming), typically 30 minutes to 1-hour post-surgery.

### Preparation of OGB solution

Acetoxymethyl (AM) ester form of Oregon Green BAPTA-1 calcium indicator (OGB-1, AM, Invitrogen, Thermo Fisher Scientific, USA) was prepared by dissolving the dye powder (50 µg) with 4µL of dimethyl sulfoxide (DMSO, Invitrogen, Thermo Fisher Scientific, USA) containing 20% of pluronic acid F-127 to create a 10 mM stock solution. The stock solution was diluted to a final concentration of 10 µM with a fresh extracellular solution (150 mM NaCl, 2.5mM KCl, and 10 mM HEPES). The solution was then filtered through a 0.2 µm sterile filter and protected from light to prevent photobleaching.

### Optical fiber recordings

Fiber photometry acquisition was performed using a custom-built system (courtesy of Prof. Arthur Konnerth, Technical University Munich; Adelsberger et al., 2005; Grienberger et al., 2012). This system utilizes a 447-nm blue-light laser diode (Osram Opto Semiconductor, Germany), which was on continuously at 40–100 μW as needed (Fig. 1A). The excitation light power at the fiber tip was calibrated using a PM100D optical power and energy meter (Thorlabs, Newton, NJ, USA). The fluorescence light path included a dichroic mirror to pass emitted green fluorescence to a photodetector (Avalanche photodiode, APD; S5343, Hamamatsu Photonics, Japan). The output signal was collected using HEKA data acquisition system (National Instruments, Austin, TX, USA), sampled at a rate of 20 or 16.7 kHz and stored in files for offline analysis.

**Figure 1.**
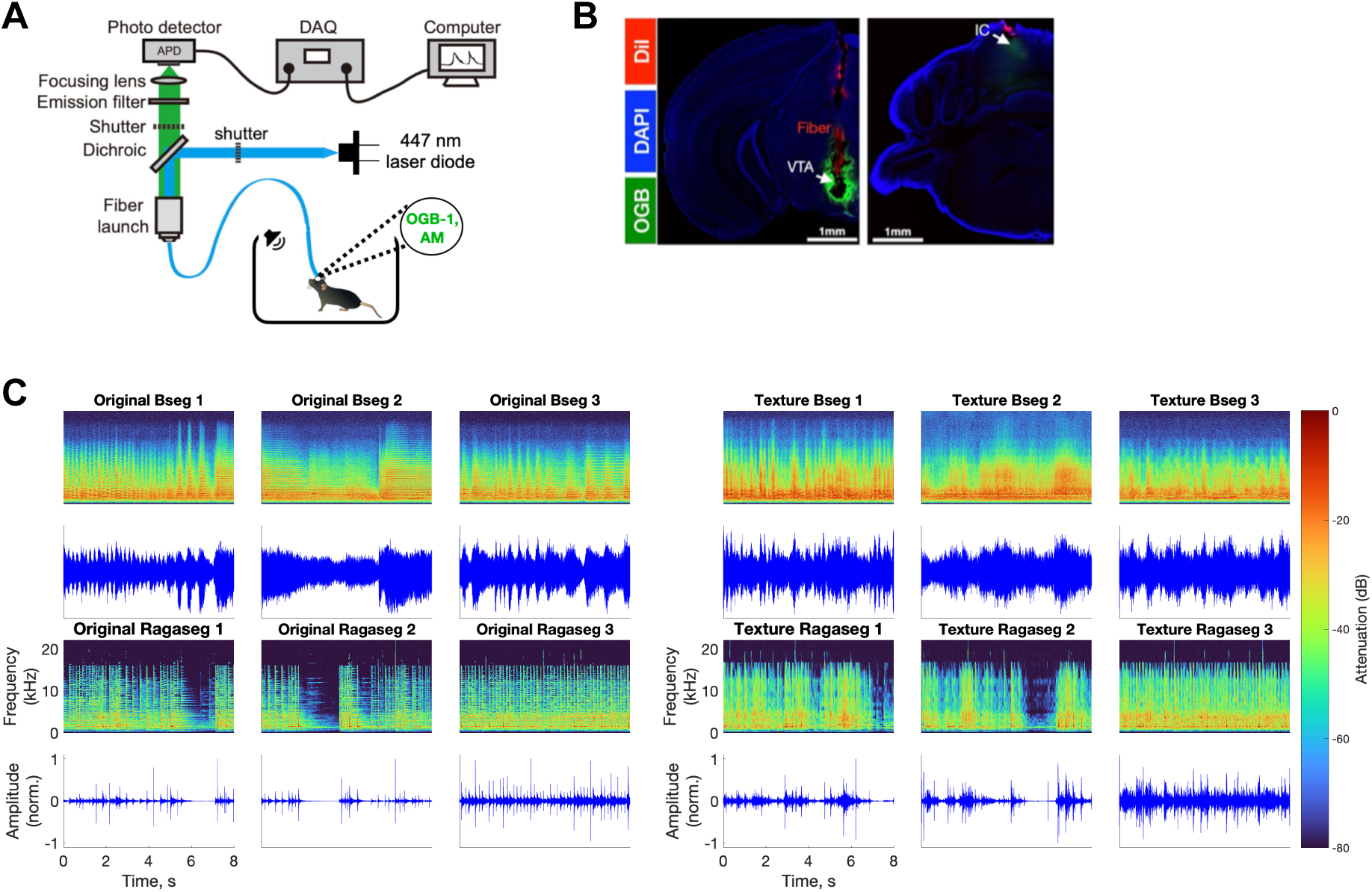
Experimental setup and complex acoustic stimuli used in the study. **A.** Fiber photometry system used for acute awake recordings in mice. **B.** Histological sections showing the fiber track (DiI, red) and OGB-1 AM (green) staining in VTA (left) and IC (right). **C.** Spectrograms and waveforms of the complex sounds: three segments from the first movement of Beethoven’s 9th Symphony and three segments of Raga music. On the right, six auditory textures were derived from the original sound segments.

### Movement tracking

We tracked the movement of the mice using a video camera (Basler AG, model acA1300-60gm, Germany) and the video recordings were analyzed by EthoVision XT (Noldus, Wageningen, Netherlands). The program tracked the center of the mouse body as it moved within the arena. In addition, the “*Distance moved*” parameter was used to quantify velocity. Timing signals recorded on the HEKA amplifier were used to synchronize the video frames with the sound presentations and the calcium signal.

### Acoustic stimuli

Simple sounds, such as broadband noise (BBN) and pure tones, were generated online using MATLAB (MathWorks) at a sampling rate of 192 kHz, converted to analog signals using a sound card (RME HDMI 9632), attenuated (PA5, TDT), and presented through a multi-field magnetic speaker (MF1, TDT). The speaker was positioned above the mouse’s open field arena. BBN and pure tones had a duration of 100 ms with 10 ms linear rise and fall times. BBN stimulation had a bandwidth of 0–50 kHz and had the same overall energy as a tone at the same sound level. BBN was presented 20 times with an inter-stimulus interval (ISI) of 1, 2 or 10 seconds. Pure tones were presented in random sequences consisting of 19 frequencies ranging from 1 to 64 kHz (3/octave) presented 10 times each, with an inter-stimulus interval (ISI) of 1, 2 or 4 seconds. BBN and tones were presented at 75 dB SPL, calibrated with a microphone at the center of the arena (Brüel & Kjær model 4189-A-021). Each such dataset is called ‘session’ below, and there could be multiple sessions recorded in the same mouse.

The complex sounds consisted of three music excerpts from the first movement of Beethoven’s Symphony No. 9 (Otto Klemperer with the Concertgebouw in Amsterdam, recorded in 1956; 8 s each), three music excerpts from classical Indian ragas (Mishra Khamaj by Pandit Shiv Kumar Sharma, Hundred Strings of Santoor, 8 s each), and six auditory textures derived from each of these sound segments (Fig. 1C). To create the auditory textures, we used the Auditory Texture Model toolbox (http://mcdermottlab.mit.edu/downloads.html). The textures share low-level statistical properties with the original excerpts, including average spectral content, amplitude modulation spectra at each frequency channel, and correlations between amplitude modulation patterns across frequency channels, but were otherwise random. These are the same sounds used in Sehrawat and Nelken (2025). The resulting 12 sounds were high-pass filtered (cutoff frequency = 1500 Hz) to remove low frequency components that are inaudible to the mice, and then presented at a sound level of 70-75 dB SPL (calibrated with the same microphone). Each sound was presented 3 or 10 times in a random sequence with an inter-stimulus interval (ISI) of 10 seconds (8 seconds of sound and 2 seconds of silence between successive presentations).

### Data analysis

All the analyses were performed on MATLAB 2021-2024 (MathWorks).

#### Fiber data processing

To correct for slow baseline drifts, the mean fluorescence over the whole trace was subtracted, and the fluorescence fluctuations were fitted with a third-degree polynomial. This polynomial fit was subtracted from the trace, and the mean fluorescence added back. The baseline was defined as the 1st percentile of this adjusted waveform (ensuring that 1% of the values were below this baseline). The fluorescence signal was then converted to ΔF/F units. Even after these corrections, slow fluctuations, likely unrelated to sensory responses, could still be present. To remove them, for the simple sounds, the data were decimated by a factor of 729 (six steps of decimation by a factor of 3), reducing the sampling rate to 23 or 28 Hz. A lower envelope was then estimated by computing the mean of the five smallest values within a symmetrical 21-sample window (spanning ±0.9 s or ±0.75 s at the decimated rate) around each time point. For the complex sounds, the data were decimated by a factor of 810 (decimation by 2, then four times by 3, then by 5), reducing the sampling rate to 21 or 25 Hz, and the lower envelope was estimated by computing the mean of the five smallest values within a symmetrical 20-second window, corresponding to 421 or 501 samples around each time point at the decimated rate. The resulting envelope traces were then smoothed using a low-pass filter with a cutoff frequency of 2.5 Hz, up-sampled back to the original sampling rate, and subtracted from the original fluorescence traces. The resulting adjusted fluorescence traces were used for all subsequent analyses.

#### Quantification of responses to broadband noise and pure tones

For some of the analyses of the responses to pure tones and BBN, we selected only trials with strong evidence for sound responses. Each trial consisted of a 200-ms pre-stimulus baseline, a 100-ms sound presentation, and 700-ms of post-stimulus activity (for a total duration of 1 s). The trials (–200 to +800 ms) were normalized between 0 (minimum) and 1 (maximum). Two single-trial measures of sensory responses were computed from the normalized trials. A responsiveness value was computed by dividing the average value of the calcium signal in the 100 ms following sound onset by the average value in the immediately preceding 100 ms of on-going activity. In addition, the rising slope of the signal was estimated as the change in fluorescence intensity during the first 50 ms after stimulus onset, and trials with a responsiveness value ≥ 1.5 and a slope ≥0.002/50 ms (in ΔF/F units) were retained for further analysis.

To quantify the fast components of these strong sound responses, we extracted the peak amplitude, peak time and response duration. The calcium peak amplitude and peak times were determined from the maximal fluorescence within a 300-ms window starting at stimulus onset. The duration of the response was estimated by the time the calcium signal decreased to half-maximum level after stimulus offset. In addition, response latency was computed from the mean of strongly responsive trials as the interval between sound onset and the time at which the fluorescence signal started changing rapidly (the inflexion point, maximum of the 2nd derivative of the fluorescence signal) in the 25 ms following sound onset.

For pure tones, a single best frequency and a single tuning bandwidth were calculated per mouse, after averaging responses across all trials pooled from multiple sessions. The best frequency was defined as the frequency eliciting the maximal mean fluorescence during sound presentations. The bandwidth was determined as the frequency range where response amplitude remained above half its maximum value, with the lower and upper bounds defined as the minimum and maximum frequencies meeting this criterion. Only the contiguous range of frequencies surrounding the best frequency was considered, with secondary peaks outside this range excluded, so the bandwidth reflects the dominant responsive frequency range even in recordings with broad or complex tuning profiles.

To check the false alarm rate of these procedures, we used on-going activity periods in which no sounds were presented. On-going activity was extracted separately for BBN and pure tone sessions. For each session, we used periods of on-going activity of approximately 10 seconds before and 10 seconds after sound presentations. The signals were segmented into consecutive 1-second windows and used as surrogate data. The surrogate data was analyzed in the same way to find strongly responsive pseudo trials for comparison with the sound-related responses.

#### Comparison of auditory responses between low- and high-velocity trials

For each trial classified as sound-responsive, we aligned the mouse instantaneous velocities, quantified as the distance traveled by the body center from one sample to the next (cm/sample, sampled at 25 samples/s, i.e., every 0.04 s), starting 200 ms before sound onset and ending at 800 ms after sound onset, and compared them with the corresponding population calcium activity. The velocity in each trial was quantified by the maximum value in the 0.8-s period following sound onset. Strongly responsive trials were divided into low- and high-velocity groups, corresponding to the lowest and highest third of velocity peak values, respectively. For each group, the mean response across trials was computed as a function of time. The effect of movement was quantified by calculating the absolute difference between the mean responses of high- and low-velocity trials at each time point. This point-by-point difference was then averaged over the trial and expressed as a percentage of the mean response in the low-velocity condition. To test the significance, a permutation test with 10,000 repetitions was performed. In each repetition, trials were randomly reassigned between low- and high-velocity groups, and the percentage difference between group means was recalculated. The p-value was the proportion of permutations where the difference was greater than or equal to the observed difference.

#### Reproducibility and discrimination

The temporal reproducibility of the trials was computed by dividing the trials into two non-overlapping halves, computing the average of each half, and correlating the two, repeating the process 1000 times and averaging the correlations. A higher correlation suggests that the trials are temporally consistent, while a lower correlation indicates variability.

To assess neuronal discrimination of the responses to music and texture sounds, responses were pooled across mice to increase trial numbers for robust neuronal discrimination analysis, and to avoid bias from unequal trial contributions. In consequence, this analysis examines population-level response patterns. Each sound had a total of 142 trials across 10 mice, with between 6 and 30 trials per sound per mouse. These were divided into two random halves (71 trials each for each sound). We then averaged the responses within each set and correlated the mean responses of the two sets for each of the twelve complex sounds. Each mean response of the first set of trials was assigned to the stimulus whose average response in the second set of trials was maximally correlated with it. This process was repeated 1,000 times to generate a confusion matrix. Accuracy was calculated with a chance level of 8% based on the use of twelve stimuli.

#### Quantification of the envelope tracking

To quantify the ability of each population to track the envelope of the complex sounds, we first determined the best frequency (BF) of each animal based on the average responses to all pure-tone trials. The sound waveforms were then filtered in a band centered on this BF, with a bandwidth defined by the equivalent rectangular bandwidth (ERB) of the auditory filter at that frequency. The filter cutoffs were set to BF − ERB/2 and BF + ERB/2. This signal served as an estimate of the effective sound that evoked the calcium responses. We then computed the magnitude of the analytic signal derived from the bandpass-filtered signal (Matlab function *hilbert*) and applied a low-pass filter with a cutoff frequency of 4 Hz to retain only the slow envelope fluctuations that are expressed in the calcium responses. For each animal and complex sound, we averaged the responses to all sound presentations, and filtered the result below 4 Hz as well. We then correlated the sound envelope with the average calcium signal (Matlab function *xcorr*). Correlation values were calculated within a lag window of 1–150 ms, and the maximum correlation within this window was used.

### Histology

At the end of the experiments, we performed histological verification of fiber placements (Fig. 1B). Mice were deeply anaesthetized with an intraperitoneal injection of pentobarbital (150 mg/kg) and transcardially perfused with PBS then by 4% paraformaldehyde (PFA) in PBS, followed by a rapid decapitation. Mouse’s heads were post-fixed in 4% PFA for 48 h at 4 °C. The brains were extracted and fixed overnight in PFA. The brains were then conserved in PBS at 4 °C. For histological determination of the injection sites and the fiber tracks, 100-μm sections were made with a vibratome (Leica VT1000). Slices were rinsed with PBS and incubated for 5 minutes with DAPI before mounting using Vectashield Mounting Medium.

## Results

We used fiber photometry to record calcium activity in the VTA of 10 freely moving mice (Fig. 1A–B) in response to broadband noise (BBN), pure tones, and complex sounds (Fig. 1C). Responses to BBN and pure tones were also obtained from the IC of 7 mice. The recordings in the IC were used for comparing the VTA responses with those recorded in a strongly auditory station using the same techniques.

### Responses to BBN

Across all trials (n = 560, 62/mouse on average, recorded from 9 mice), most of the trials exhibited a sustained increase in calcium signal (Fig. 2A). In a minority of trials (∼10%), the calcium signal decreased during stimulus time. The subset of strongly responsive trials (n = 197, 35%; see Methods for selection criteria) exhibited a pronounced time-locked calcium response (Fig. 2B). The average response profile for these trials had a rapid rise in ΔF/F with a latency of 15 ms peaking within approximately 100 ms after stimulus onset, followed by a gradual decay (Fig. 2C). To verify that the observed responses were stimulus-driven rather than spontaneous fluctuations, we applied identical selection criteria to pseudo trials extracted from periods of on-going activity. Only 90 of 696 pseudo trials (13%) met the strongly responsive criteria, significantly fewer than during sound stimulation (Fisher’s exact test, p < 0.0001, Fig. 2D–E), and their mean response was significantly smaller than the mean response of strongly responsive trials measured over the 200 ms response window following sound offset (hereafter referred to as the post-offset response window; Wilcoxon rank-sum test, p = 0.018). This window excluded the 100 ms period during sound presentation, which was used to select strongly and strongly responsive pseudo trials. Furthermore, the studentized (normalized by its standard error) mean response of strongly responsive pseudo trials was substantially smaller (∼3) compared with strongly responsive trials (∼14) across the response window (Fig. 2F).

**Figure 2.**
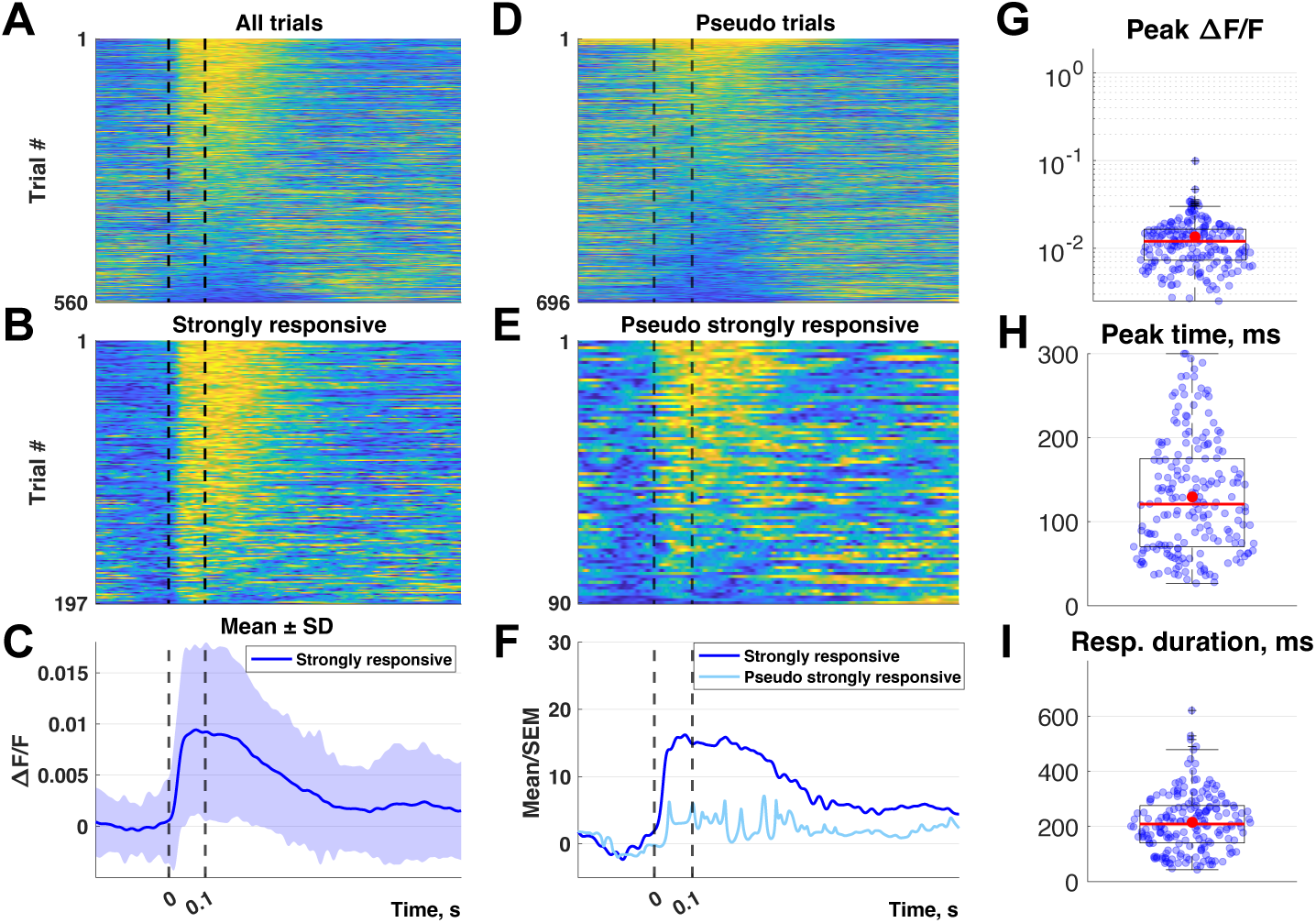
VTA responses to broadband noise. **A.** All normalized trials showing VTA responses to broadband noise (BBN). Trials are ordered according to the maximum average calcium level within 300 ms after sound onset (black dashed lines indicate sound onset and offset). **B.** Subset of strongly responsive trials, ordered as in (A). **C.** Mean calcium response to BBN (± SD) for strongly responsive trials. **D–E.** Activity patterns during spontaneous activity periods. (D) Full set of pseudo trials showing spontaneous activity, ordered as in (A). (E) Subset of strongly responsive pseudo trials, ordered as in (A). **F.** Time course of the studentized (Mean/SEM) mean of the strongly responsive trials (dark blue) and strongly responsive pseudo trials (light blue). **G-I.** Box plots summarizing response parameters for strongly responsive trials: calcium peak amplitude (G), peak time (H), and response duration (I). Red dots indicate mean values for each parameter.

Figures 2G–I summarize the single-trial calcium peak amplitudes, peak times, and response durations for the strongly responsive trials. Since the decay time constants of the calcium indicator are longer than the duration of the BBN stimuli, the calcium signal largely represents the time integral of the neural activity evoked by the stimulus. Therefore, late peak times likely indicate a sustained, tonic response, while earlier peak times likely indicate phasic responses. The strongly responsive trials had fast calcium increases with a median peak amplitude of 0.010 ΔF/F (Fig. 2G) and peak times centered around 120 ms (Fig. 2H), suggesting sustained responses throughout stimulus presentation time. The response duration extended for a hundred ms after sound offset (median ≈ 210 ms; Fig. 2I).

### Responses to pure tones

Population activity recorded in response to pure tones was first averaged across all presentations of each frequency in each animal. Representative examples from three mice derived from the full set of trials, showing responses at specific frequencies are shown in Fig. 3A. The strongest responses were almost always evoked by tones in the most sensitive part of the hearing range of the C57BL6 mice, between 4 and 16 kHz (Willott, 1986; Willott, 1993; Bowen, 2020). The distributions of BF and bandwidths across animals (Figs. 3B–C) showed that most recordings were best driven by low to mid frequencies (<16 kHz) and exhibited frequency tuning between 2 to 6 octaves.

**Figure 3.**
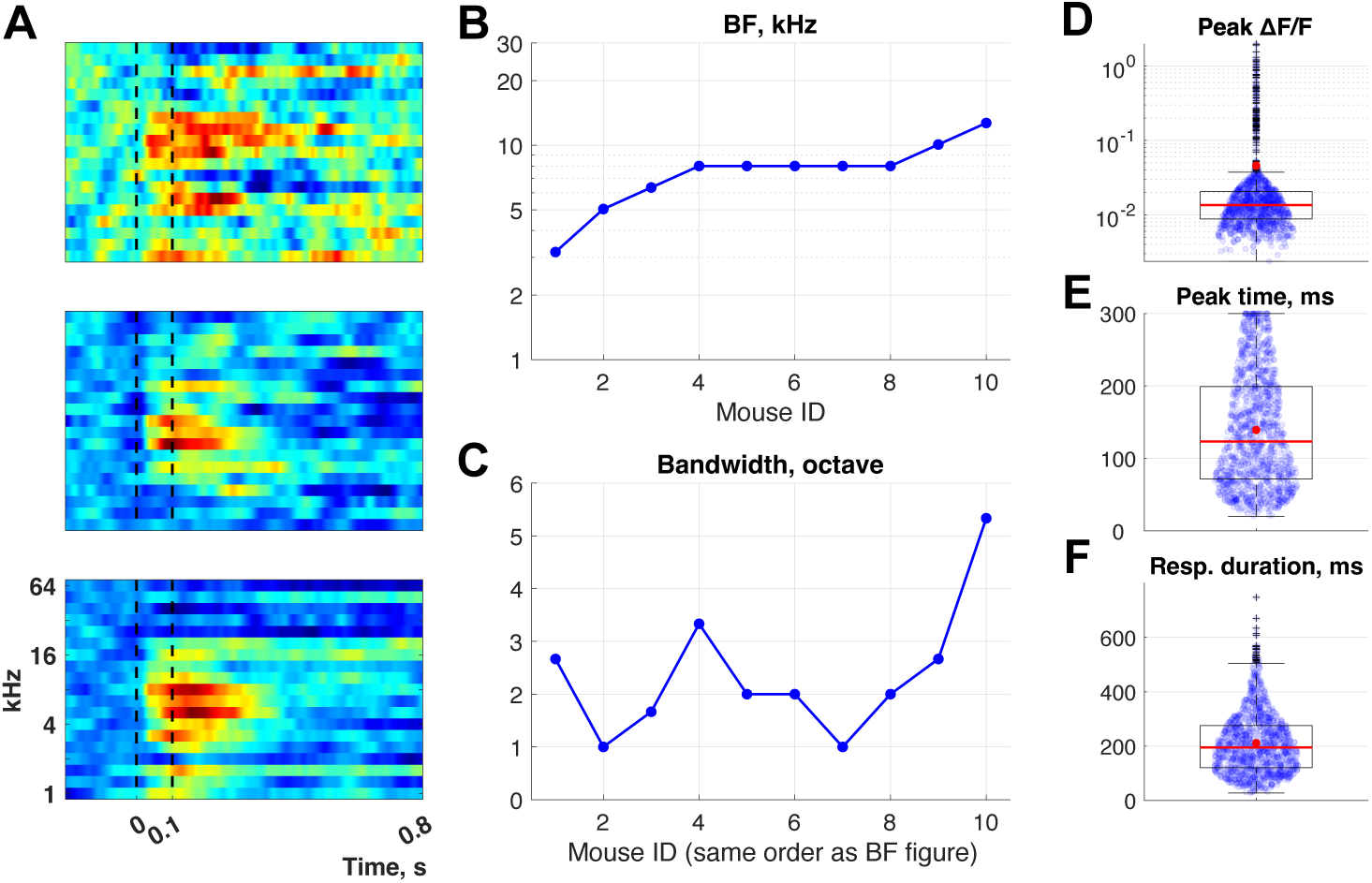
VTA responses to pure tones. **A.** Representative examples of population activity in response to pure tone stimuli (1 to 64 kHz, 100 ms, black dashed lines) from three mice. **B-C.** Best frequency (B) and bandwidth (C) for each mouse, ordered by the estimated best frequency. **D-F.** Boxplots showing calcium peak amplitudes from strongly responsive trials across all tested frequencies (1–64 kHz), along with corresponding peak times, and response durations. Red dots indicate mean values for each parameter.

For analyzing single trials, we extracted the subset of strongly responsive trials (1181 of 6460, 18.3%). As for BBN, when the same criteria were applied to pseudo trials extracted from periods of on-going activity, significantly fewer trials were classified as strongly responsive (105 of 819, 13%; Fisher’s exact test, p<0.0001, Supplementary Fig. 1), and their mean response was significantly smaller than the mean response of strongly responsive trials in the post-offset response window (Wilcoxon rank-sum test, p < 0.001). The studentized mean response of the strongly responsive pseudo trials was smaller (∼2) compared with strongly responsive trials (∼7) across the response window (Supplementary Fig. 1).

Figures 3D–F summarize the distributions of calcium peak amplitudes, peak times, and response durations for strongly responsive trials (collapsed across all frequencies). Similar to the broadband noise responses, strongly responsive trials had fast rise time (latency of the mean response: 13 ms), with a median peak amplitude of 0.013 ΔF/F (Fig. 3D) and peak times around ∼120 ms (Fig. 3E). The average response duration extended for about a hundred ms after stimulus offset (median ≈ 190 ms; Fig. 3F).

### Movements only weakly affected the responses

The recordings were performed while the animals were freely-moving. We therefore checked whether the strongly responsive trials were influenced by movements that co-occurred with the sound presentations, potentially causing measurement artifacts (Fig. 4). Velocities during strongly responsive trials to both BBN and pure tones were extracted (see Methods; Fig. 4A). We studied the relationships between movement and calcium signals using the peak calcium response amplitude within a 300-ms window from stimulus onset and the peak velocity values within 0.8 seconds from stimulus onset (Fig. 4B). The effect of velocity magnitude on peak responses was assessed using a linear mixed-effects model for log-transformed peak responses (fixed effects: log(max velocity) and stimulus type (BBN or pure tones) and their interaction; random intercepts and slopes for animals). All main effects were non-significant (log(max velocity): F(1,1173) = 0.088, p = 0.77; stimulus type: F(1,1173) = 0.014, p = 0.90; interaction: F(1,1173) = 0.019, p = 0.89).

**Figure 4.**
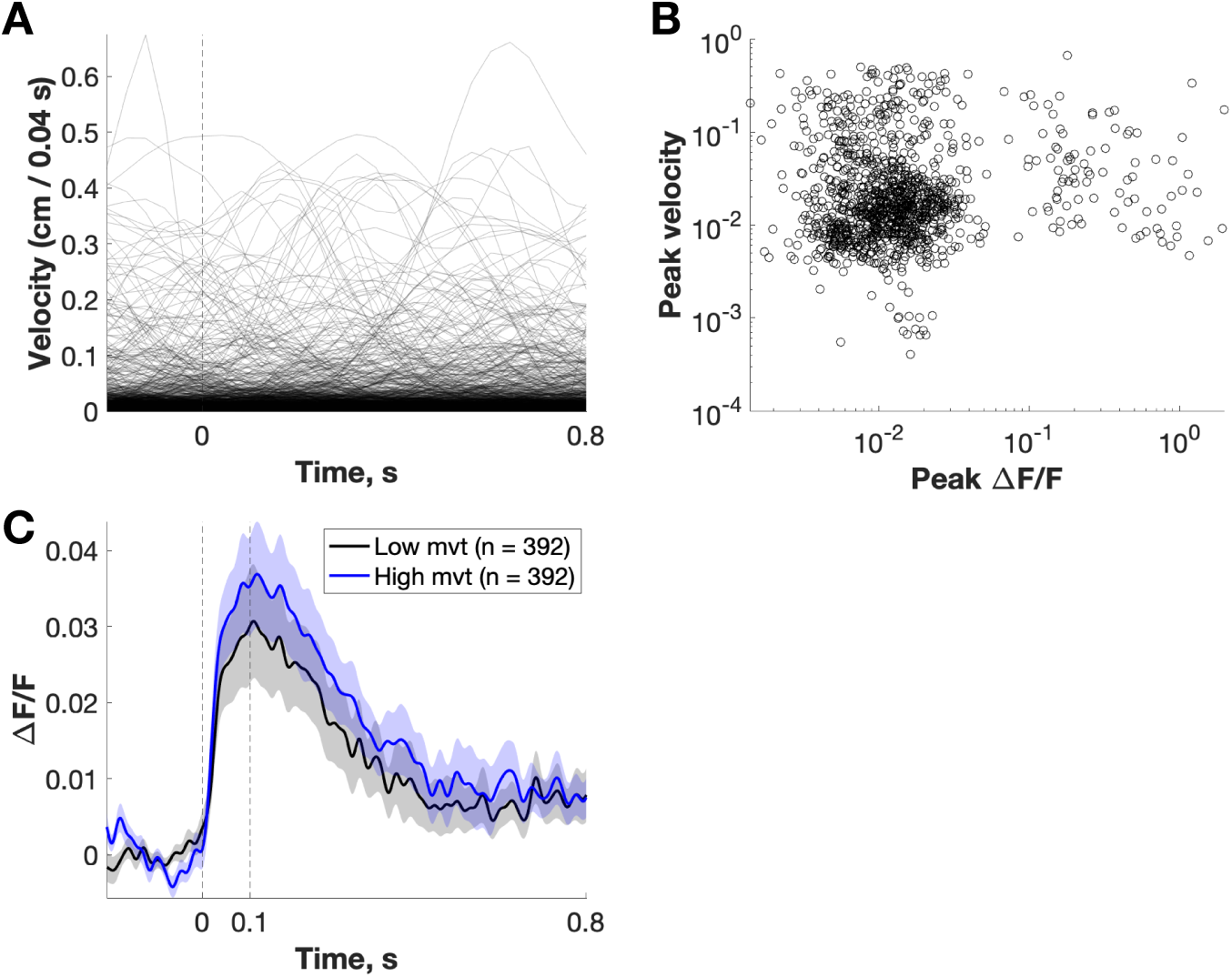
Body movements weakly influenced responses in the VTA. **A.** Velocity (cm/0.04 s) during 1-s trial presentations. Data include all strongly responsive trials to both broadband noise and pure tone stimuli. **B.** Scatter plot of peak velocity as a function of peak calcium responses for strongly responsive trials. **C.** Mean calcium responses (ΔF/F) aligned to stimulus onset (vertical dashed line at 0ms) for the 1/3 of trials with lowest velocity and the 1/3 with highest velocity. Shaded areas represent ± SEM.

To illustrate the weak effect of movement on the calcium responses, we formed low- and high-velocity groups (Fig. 4C). The low-velocity group consisted of the one third of the strongly-responsive trials with the lowest velocity peak values, while the high-velocity group included the one third of the strongly-responsive trials with the highest velocity peak values. The average calcium responses in the two subsets of trials are displayed in Fig. 4C. The responses in the high velocity trials were slightly larger than in the low velocity trials. We calculated the percentage difference over the entire 1-second trace as the mean absolute point-by-point difference between high- and low-velocity trials, normalized by the mean response during low-velocity periods. The responses showed a small increase (∼33%) during high-velocity periods compared to low-velocity periods, but this difference was not statistically significant (permutation test, p = 0.46). Together, these analyses suggest that movement had at most a weak influence on the sensory responses in the VTA.

### Sound-evoked calcium dynamics in the inferior colliculus

We compared the VTA responses with those recorded in a major auditory area, the inferior colliculus. We found reliable sound-evoked responses to BBN (n = 320 trials, Fig. 5A), as expected for this midbrain auditory center. Even though BBN is known to drive well population responses in the IC, only about half of the trials (n = 159, 50%, Fig. 5B-C) were classified as strongly responsive (see Methods for selection criteria). When the same selection criteria were applied to pseudo trials extracted from periods of on-going activity, only a small fraction of trials were classified as strongly responsive (28 of 308, 9%), significantly fewer than during sound-evoked trials (Fisher’s exact test, p < 0.001, Fig. 5D-F). Their mean response was also significantly lower than that of strongly responsive sound-evoked trials in the post-offset response window (Wilcoxon rank-sum test, p < 0.0001) and the studentized values were ∼4-fold lower for strongly responsive pseudo trials (mean ∼5) compared with strongly responsive trials (mean ∼18) across the response window (Fig. 5F). Strongly responsive trials showed pronounced peak ΔF/F amplitudes (median: ∼0.011, Fig. 5G) with rapid-onset activity (latency of the mean response: 15 ms) reaching their maximum at ∼110 ms (median, Fig. 5H) and decaying over ∼190 ms (median, Fig. 5I). As in the VTA, about 10% of the trials exhibited decreases in calcium signals. The proportion of strongly responsive trials was significantly lower in the VTA than in the IC (35% vs. 50%; Fisher’s exact test, p < 0.0001).

**Figure 5.**
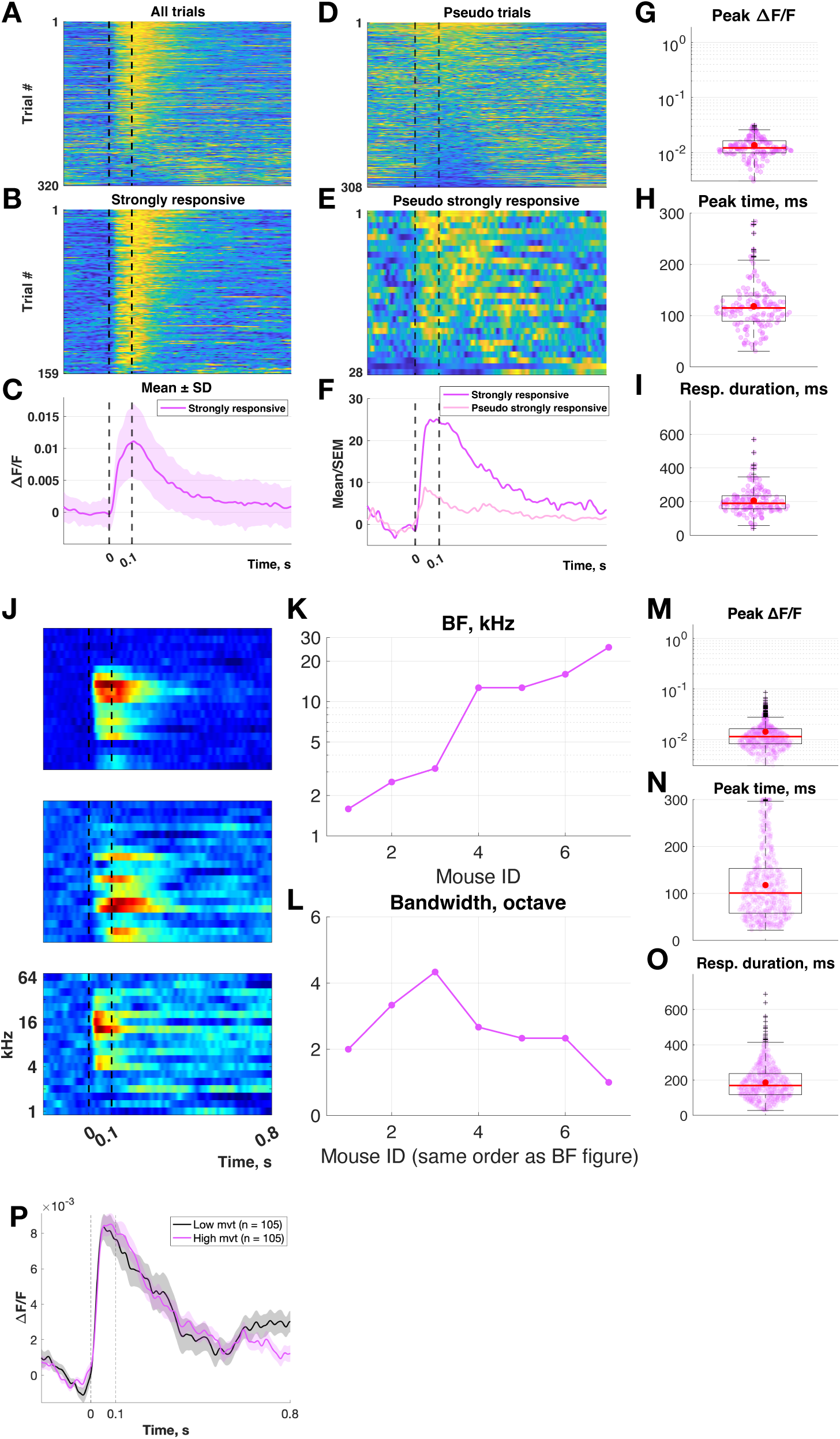
Sound-evoked calcium responses and movement-related modulation in the awake inferior colliculus. **A.** All normalized trials showing IC responses to broadband noise. Trials are ordered according to the maximum average calcium level within 300 ms after sound onset (black dashed lines indicate sound onset and offset). **B.** Subset of strongly responsive trials ordered as in (A). **C.** Mean response to BBN (± SD) for strongly responsive trials. **D-F.** All pseudo trials from periods of spontaneous activity in the absence of sound. (D): all pseudo trials ordered according to maximum activity within the analysis window of 300 ms. (E): the subset of strongly responsive pseudo trials ordered as in the top panel. (F): studentized mean of the strongly responsive trials (dark magenta) and pseudo trials (light magenta). **G–I.** Boxplots showing, respectively, calcium peak amplitudes in strongly responsive trials, peak times, and response durations. Red dots show the mean values obtained for each parameter. **J.** Average population activity in response to pure tone stimuli (1 to 64 kHz, 100 ms; indicated by black dashed lines) from three mice. **K-L.** Best frequency (K) and bandwidth (L) for each mouse, ordered by the estimated best frequency. **M-O**. Boxplots showing calcium peak amplitudes from strongly responsive trials across all tested frequencies (1–64 kHz), the corresponding peak times, and response durations. Red dots show the mean values obtained for each parameter. **P.** Mean calcium responses (ΔF/F) aligned to stimulus onset (vertical dashed line at 0ms) for the 1/3 of trials with lowest velocity and the 1/3 with velocity values. Shaded areas represent ± SEM.

To assess differences in the properties of strongly responsive trials, we used linear mixed-effect models. The models had recording site (VTA or IC) as a fixed factor and random intercepts and slopes for animals within each site. There were no significant differences between VTA and IC in any of the parameters of the strongly responding trials (peak responses: F(1,354) = 0.92, p = 0.34; peak times: F(1,354) = 0.07, p = 0.79; response durations: F(1,354) = 0.04, p = 0.84).

Pure tone stimuli (1–64 kHz) evoked clear tuning (Fig. 5J) with best frequencies ranged from 1.5–25 kHz and bandwidths between 1–4 octaves (Figs. 5K-L). To check whether the distribution of BFs and bandwidths were different in the two recording sites, we used a permutation test. Animals were shuffled between the VTA and IC, preserving group sizes, and the distributions of BFs and bandwidths were compared using a Kolmogorov-Smirnov test. Neither BF distribution nor bandwidth distribution differed between the two recording sites (BFs: KS = 0.471, p = 0.15; Bandwidth: KS=0.457, p=0.17).

Across frequencies, responses reached a median peak ΔF/F of 0.009 at ∼100 ms, lasted ∼170 ms, and showed a mean latency of 13 ms (Figs. 5M–O). Pseudo trials extracted from on-going activity yielded significantly fewer strongly responsive trials than evoked activity (10.0%, 31/328 vs. 22.2%, 718/3230; Fisher’s exact test, p < 0.0001; Supplementary Fig. 2). Moreover, the mean response was significantly lower than that of strongly responsive sound-evoked trials during the post-offset response window (Wilcoxon rank-sum test, p<0.0001) and the studentized values were markedly lower for strongly responsive pseudo trials (mean ∼2) compared with strongly responsive trials (mean ∼20) across the response window (Supplementary Fig. 2). Pure-tone responsive trials showed several differences between VTA and IC. While peak response amplitudes were not significantly different between VTA and IC (ANOVA: F(1,1897) = 1.86, p = 0.17), peak response times occurred significantly later in VTA (average: ∼140 ms vs. ∼120 ms; ANOVA: F(1,1897) = 3.98, p = 0.046), and response durations were significantly longer in VTA than in IC (average: ∼ 210 ms vs. ∼190 ms; ANOVA: F(1,1897) = 4.06, p = 0.044).

Movement effects on the responses were also similar in IC and VTA (Fig. 5P). The effect of movement magnitude on peak responses was assessed using a linear mixed-effects model for log-transformed peak responses (fixed effects: log(max movement) and stimulus type (BBN or pure tones) and their interaction; random intercepts and slopes for animals). Both main effects and the interaction were non-significant (log(max movement): F(1,313) = 1.45, p = 0.23; stimulus type: F(1,313) = 2.86, p = 0.09; log(max movement) × stimulus: F(1,313) 1.30, p = 0.26). There was a slight average decrease (19%) in sound-responsive trial responses (including both BBN and pure tones) during high-movement periods compared to low-movement periods (Fig. 5P), but this difference was not statistically significant (permutation test, p = 0.67). In summary, movement effects on sound-evoked responses were modest and not statistically significant.

### Responses to complex sounds

With stimuli spanning 8s, measures such as peak amplitude and latency do not provide sufficiently detailed characterization of the responses. We therefore quantified temporal reproducibility, envelope tracking, and the discriminability of the responses to the different complex stimuli.

We lowpass-filtered (<4 Hz) the responses to the complex sounds in order to enhance those response components that are most likely representing envelope locking. The temporal response profiles to the different sounds varied, with some sounds eliciting more pronounced transients than others (Fig. 6A). To quantify temporal reproducibility of the single trials, we computed inter-trial correlations (ITC) using responses to the repetitions of each stimulus (Fig. 6B). Averaged ITC values varied across stimuli, with some stimuli eliciting consistent responses (positive ITC) and others not. Average ITC values ranged from -0.31 to 0.42 and were positive for 10/12 of the sounds. Note that tR3 differed from the other stimuli, as its average ITC was negative. The average decoding accuracy (see Methods) reached 19% (Fig. 6C), although individual stimuli had decoding accuracy as high as 60%. This was above chance level (8%), suggesting that VTA activity carries a detectable amount of information about the acoustic structure of the stimuli. Temporal reproducibility and stimulus discriminability were related: sounds with high ITC were also those that showed higher probability of being correctly classified (Pearson r = 0.79, p = 0.002; Fig. 6D).

**Figure 6.**
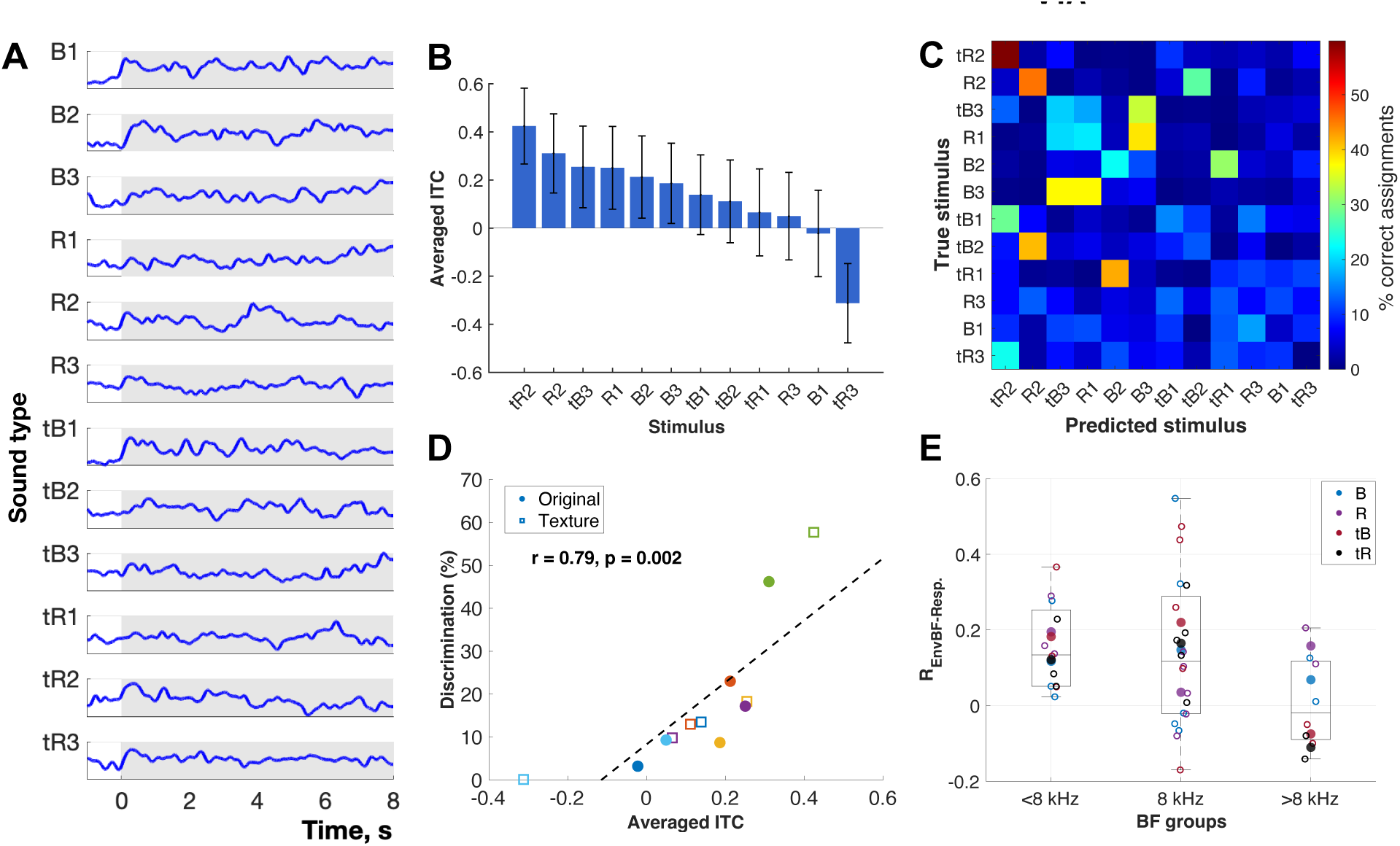
Temporal reproducibility and information content of VTA responses to complex sounds. **A.** Average (<4 Hz) VTA responses evoked by each complex sound stimulus, low-passed filtered at 4 Hz. Responses were normalized in a 0-1 scale across all stimuli together. Sound categories include three excerpts from the first movement of Beethoven’s 9th Symphony (B1–B3), three excerpts of Raga music (R1–R3), and auditory textures derived from each (tB1–tB3; tR1–tR3). **B.** Temporal reproducibility across trials, quantified using inter-trial correlation (ITC). Stimuli are ranked in descending order of mean ITC values. **C.** Confusion matrix showing classification accuracy for discriminating the 12 stimuli from neural responses. Stimuli are ranked in the descending order of mean ITC values. **D.** Relationship between ITC and classification accuracy for original (filled circles) and texture-derived (open squares) stimuli. Each color represents a stimulus and its texture: B1/tB1 (blue), B2/tB2 (red), B3/tB3 (yellow), R1/tR1 (purple), R2/tR2 (green), and R3/tR3 (cyan). **E.** Correlation values between the mean responses and the sound envelopes filtered around the best frequency (BF) of each mouse. Data are grouped into three BF groups: <8kHz (n=3 mice), 8kHz (n=5 mice) and >8kHz (n=2 mice). Filled points indicate the mean correlation averaged across mice for each sound type, while open circles represent individual mice: B: Beethoven excerpts (blue), R: Raga excerpts (purple), tB: Beethoven texture excerpts (dark red), tR: Raga texture excerpts (black).

Finally, we assessed the envelope representation of the responses to complex sounds. Supplementary Fig. 3A presents the envelopes of the frequency band around 8 kHz for all complex sounds. Supplementary figure 3B displays the envelopes of each frequency band of the B1 sound. While the overall time course of the envelopes was similar across frequency bands, subtle changes were present as center frequency shifts from low to high frequencies.

We compared the temporal response pattern to each of the complex sounds with the envelope of the frequency band corresponding to the BF of each animal (Fig. 6E). To assess the effects of BF, animals were grouped into three BF categories (<8kHz: n=3 mice, 8kHz: n=5 mice and >8kHz: n=2 mice). We initially fitted a linear mixed-effects model with correlation values as the dependent variable, stimulus type (Beethoven, Raga, Beethoven textures, Raga textures), BF group, and their interaction as fixed effects. The interaction between stimulus type and BF group was not significant (F(6,28) = 2.42, p = 0.21), and removing the interaction did not significantly reduce the goodness of fit (likelihood ratio test: χ²(6) = 8.20, p = 0.22). Therefore, we used a model with the main effects of stimulus type and BF group only.

The model indicated that the intercept (accounting for the mean correlation across all BFs and stimuli) was significantly different from zero (F(1,34) = 6.35, p = 0.017). No significant main effects of BF group (F(2,34) = 2.43, p = 0.104) or stimulus type (F(3,34) = 0.21, p = 0.887) were observed. Overall, these results indicate that correlation between VTA activity and the temporal envelope of the stimuli was weak and did not differ across BF groups or stimulus types.

## Discussion

In awake, freely moving mice, VTA populations responded to a wide range of sounds—including broadband noise, pure tones, and complex stimuli, demonstrating that the VTA is sensitive to auditory input. When responses occurred, a significant fraction of trials had a clear rapid sound-related response (Figs. 2 and 5, and comparison with the pseudo trials). Strongly responsive trials to both BBN and pure tone stimuli were remarkably similar to strongly responsive trials to the same stimuli in IC. Population responses to complex sounds showed a weak, but significant, envelope tracking and trial-to-trial temporal reproducibility. This resulted in a significant, even if low, discrimination performance.

### VTA neurons respond to many sounds

Few studies have investigated how VTA respond to sensory stimuli. Here, we measured calcium activity from populations of VTA neurons. The most important finding of this paper is the observation of reliable auditory-evoked responses, in spite of the lack of behavioral relevance of the sounds. This suggests that at least some neurons in the VTA respond to auditory inputs independently of their behavioral meaning. The VTA is heterogenous, with dopaminergic (∼60%), GABAergic (∼35%), and glutamatergic (∼5%) neurons (Yamaguchi et al., 2015). It could be that just one of these subsets of neurons is unconditionally sound-responsive. Such activity is not unprecedented: electrophysiological recordings in awake, freely moving cats have shown that VTA dopaminergic neurons respond to unconditioned auditory stimuli, such as clicks at 73 dB SPL (Horvitz et al., 1997). Consistent with this, Wei et al. (2024) reported auditory responses in VTA neurons of awake, head-fixed naïve mice using in vivo electrophysiology. In their study, broadband white noise (2–64 kHz) was presented at intensities ranging from 0 to 100 dB SPL, and VTA neurons exhibited response latencies of 10–15 ms (mean ∼12 ms), with an average response threshold around 60 dB SPL.

The auditory responses we document here are similar to those recorded with the same methods in the inferior colliculus. The averaged strongly-responsive trials in the VTA exhibited latencies of ∼13–15 ms, comparable to those in the IC. While recent studies in mice reported evoked responses to loud broadband sounds, our results extend this observation to responses evoked by a broader range of auditory stimuli.

Engelhard et al. (2019) recorded single dopaminergic neurons in the VTA and found response components which were not directly related to reward but rather to behaviorally-relevant variables (spatial location, kinematics and behavioral choices). Their results are in line with our results showing that VTA responses may be reliably activated by sensory events which are not associated with reward. We found that the frequency range over which VTA responded to pure tones was similar to IC (Fig. 3A, 5H-I). In the IC, the broad population-level frequency range likely emerges from the integration of calcium signals from many neurons, each of which had narrow tuning curves. The VTA’s broad tuning could arise from similar population-level integration of narrowly tuned neurons, but also from different mechanisms, such as the summation of responses from individual neurons that are themselves more broadly tuned. In fact, the population involved in auditory processing in the VTA is likely smaller than in the IC, since we do not expect all VTA neurons to show auditory sensitivity. This may contribute to the weaker responses we observed to pure tones.

Finally, using complex, rich sounds, we found that trial-to-trial reproducibility was generally low, and population responses tracked sound envelopes only weakly, resulting in poor discrimination of complex stimuli.

### Auditory inputs to the VTA

To the best of our knowledge, there is no direct connection from the auditory pathway to VTA. However, VTA neurons receive inputs from numerous brain regions, with a total of at least 29 identified sources (Soden et al., 2020; Watabe-Uchida et al., 2012). Many of these inputs show auditory responses, including the orbitofrontal cortex (Winkowski et al., 2018), the medial prefrontal cortex (Zhao et al., 2019), the dorsal striatum (Bordi and LeDoux, 1992), the dorsal raphe nucleus (Waterhouse et al., 2004) and the locus coeruleus (Hervé-Minvielle and Sara, 1995).

It could be that the VTA receives auditory information from the dorsal raphe nucleus (DRN). The DRN receives direct auditory inputs from the inferior colliculus (Pollak-Dorocic et al., 2014), and there are strong reciprocal connections between the VTA and the DRN nuclei. Auditory-evoked responses have been found in the DRN to simple stimuli (clicks and tones; Waterhouse et al., 2004) and to vocalizations (Heym et al., 1982; Rasmussen et al., 1986).

A new non-canonical auditory pathway has been described recently by Zhang and colleagues (2020). It originated from the cochlear nucleus and, through the caudal pontine reticular nucleus (PRN), pontine central gray (PCG), and medial septum (MS), extended to the entorhinal cortex (EC). PCG projects to VTA, and could underlie at least some of the VTA auditory sensitivity.

However, the neurons along this pathway responded primarily to high-level white noise, while our results showed responses to pure tones as well. Thus, this reticular-limbic pathway may not be solely responsible for the observed auditory responses in VTA. Indeed, more recently, the same team discovered that while silencing PCG, VTA neurons showed a decrease in activity but still responded to noise (Wei et al., 2024), and here we show that response latencies in the VTA were similar to those observed in the inferior colliculus. These observations suggest the presence of an additional, PCG-and DRN-independent pathway from the auditory system to the VTA.

### Functional implications

The wide and diverse input connectivity of the VTA potentially contributes to the heterogeneous evoked activity we recorded, particularly for complex sounds, reflecting the integration of sensory, cognitive, and motivational information. In parallel, the VTA sends widespread projections to regions involved in reward processing, sensory integration, emotional regulation, and behavioral control, including the nucleus accumbens, amygdala, cortex, hippocampus, ventral pallidum, periaqueductal gray, bed nucleus of the stria terminalis, olfactory tubercle, and locus coeruleus (Morales and Margolis, 2017). Through this broad connectivity, VTA activity is well positioned to modulate downstream circuits in a context-dependent manner.

Sound-evoked activity in the VTA may contribute to shaping perception and behavior by linking auditory input to motivational and contextual representations. VTA activity is believed to play a critical role in plasticity, influencing attention, learning, and memory, largely mediated by dopamine (Gittelman et al., 2013; Happel, 2016), reinforcing the view of the VTA as a multifunctional hub beyond its role as a reward-processing center. Within this framework, our results suggest that acoustic signals in the VTA may be transformed, highlighting their potential to convey distinct emotional or motivational significance.

### Limitations of the study

The key limitation of this study is that our recordings were performed using fiber photometry, which reflects the activity of populations of neurons rather than single-neuron activity. Our techniques made it impossible to record subtype-specific signals – our calcium signals reflect the combined spiking activity of the dopaminergic, GABAergic and glutamatergic neurons in the VTA.

## Additional information section

### CRediT authorship contribution statement

**Samira Souffi:** Investigation, Data curation, Formal analysis, Methodology, Validation, Resources, Writing – original draft, Writing – review & editing.

**Israel Nelken:** Conceptualization, Supervision, Project administration, Funding acquisition, Methodology, Validation, Writing – review & editing.

## Acknowledgements

We thank Dr. Dina Moshitch and Dr. Kamini Sehrawat for technical assistance with the fiber photometry system. IN holds the Milton and Brindell Gottlieb Chair in Brain Sciences.

---

**Supplementary figure 1.**
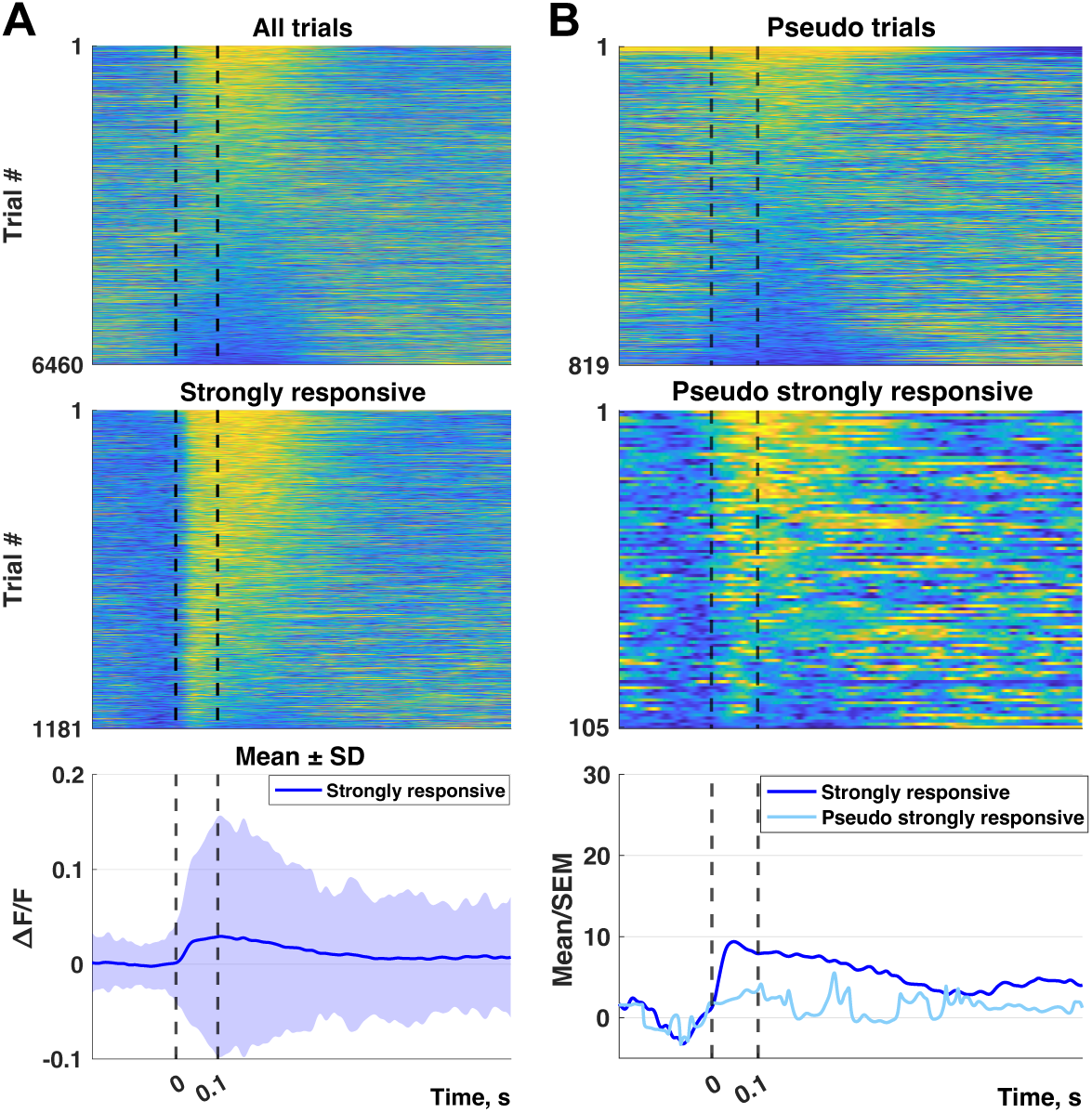
Activity patterns during evoked and spontaneous activity periods in VTA during pure tone sessions. **A.** All normalized trials (n=6460) showing VTA calcium responses to pure tones, ordered by maximum average activity within 300 ms after sound onset (dashed lines indicate sound onset and offset). Middle: subset of strongly responsive trials (n=1181), ordered as above. Bottom: mean calcium response (ΔF/F ± SD) for the strongly responsive trials. **B.** All pseudo trials (n=819) extracted from spontaneous activity periods, ordered as in (A). Middle: subset of strongly responsive pseudo trials (n=105), ordered as in (A). Bottom: time course of the studentized mean of the strongly responsive trials (dark blue) and strongly responsive pseudo trials (light blue).

**Supplementary figure 2.**
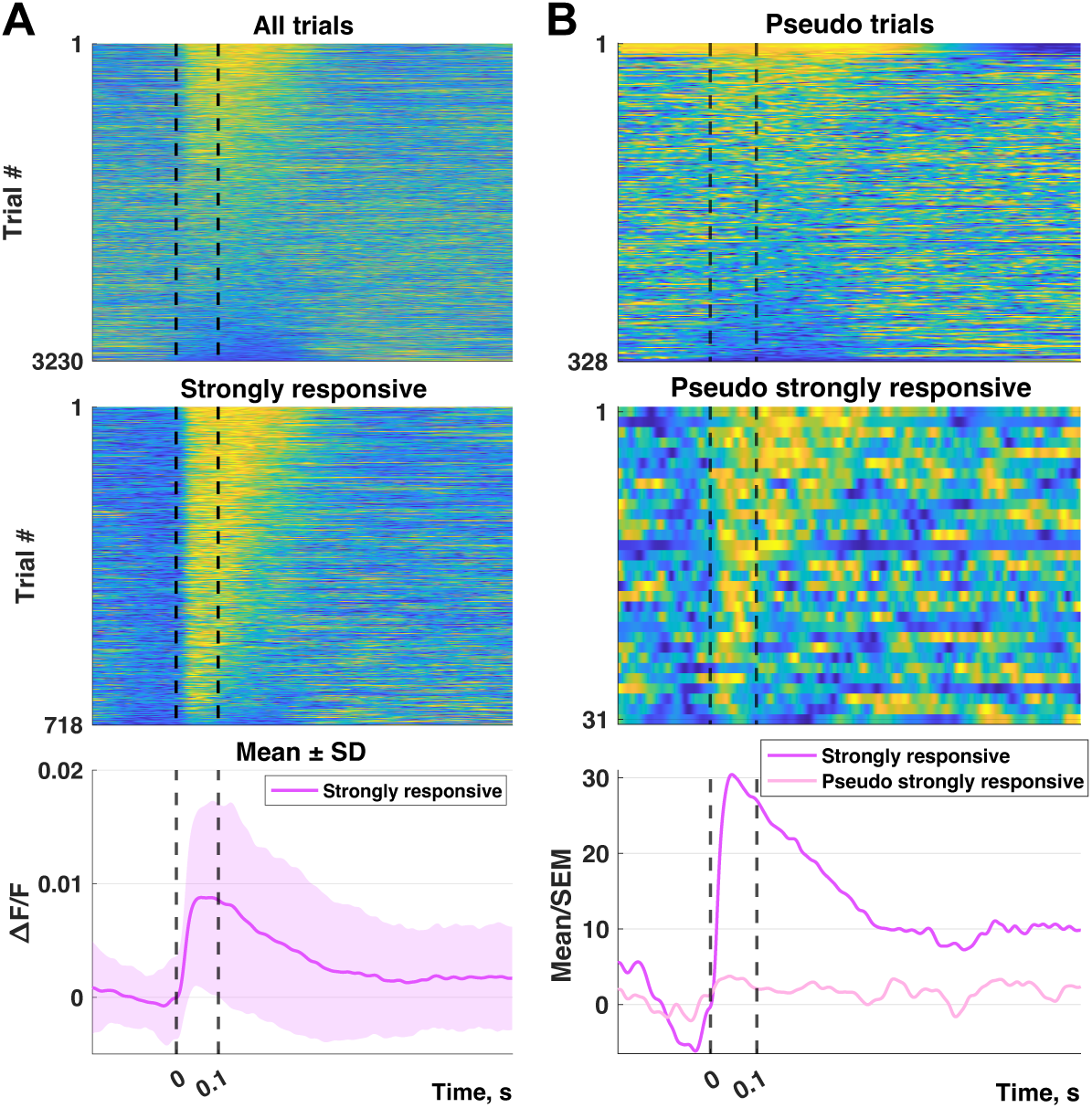
Activity patterns during evoked and spontaneous activity periods in IC during pure tone sessions. **A.** All normalized trials (n=3230) showing IC calcium responses to pure tones, ordered by maximum average activity within 300 ms after sound onset (dashed lines indicate sound onset and offset). Middle: subset of strongly responsive trials (n=718), ordered as above. Bottom: mean calcium response (ΔF/F ± SD) for strongly responsive trials. **B.** All pseudo trials (n=328) extracted from spontaneous activity periods, ordered as in (A). Middle: subset of strongly responsive pseudo trials (n=31), ordered as in (A). Bottom: time course of the studentized mean of the strongly responsive trials (dark blue) and strongly responsive pseudo trials (light blue).

**Supplementary figure 3.**
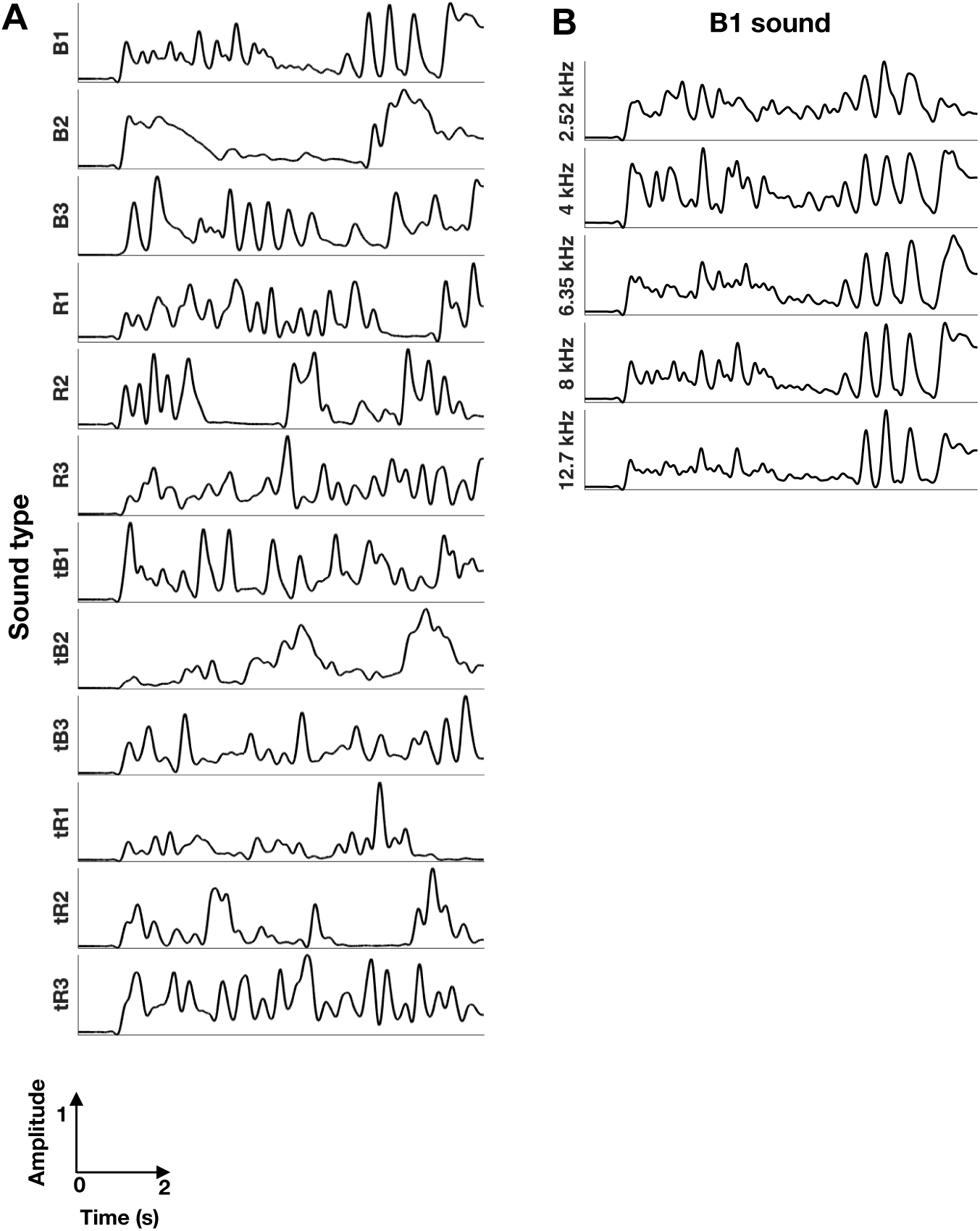
Envelopes of complex sounds. **A.** Envelopes of the 1-octave band centered at 8 kHz, low pass filtered below 4 Hz. **B.** The sound envelope of sound B1, bandpass filtered at different central frequencies. The envelopes were then low pass filtered below 4 Hz.

